# Genome-wide Chromatin Accessibility is Restricted by ANP32E

**DOI:** 10.1101/2020.09.08.288241

**Authors:** Kristin E. Murphy, Fanju W. Meng, Claire E. Makowski, Patrick J. Murphy

## Abstract

Genome-wide chromatin state underlies gene expression potential and cellular function. Epigenetic features and nucleosome positioning contribute to the accessibility of DNA, but widespread regulators of chromatin state are largely unknown. Our study investigates how control of genomic H2A.Z localization by ANP32E contributes to chromatin state in mouse fibroblasts. We define H2A.Z as a universal chromatin accessibility factor, and demonstrate that through antagonism of H2A.Z, ANP32E restricts genome-wide DNA access. In the absence of ANP32E, H2A.Z accumulates at promoters in a hierarchical manner. H2A.Z initially localizes downstream of the transcription start site, and if H2A.Z is already present downstream, additional H2A.Z accumulates upstream. This hierarchical H2A.Z accumulation coincides with improved nucleosome positioning, heightened transcription factor binding, and increased expression of neighboring genes. Thus, ANP32E dramatically influences genome-wide chromatin accessibility through refinement of H2A.Z patterns, providing a means to reprogram chromatin state and to hone gene expression levels.

## INTRODUCTION

Access to specific regions of DNA by transcription factors and polymerase machinery is critical for the regulation of gene expression. The packaging of DNA, including nucleosome occupancy, positioning, wrapping, and the presence of chromatin modifications, termed ‘epigenetic marks’, has been implicated in regulating DNA accessibility^1^. However, the degree to which these epigenetic marks regulate transcription factor (TF) binding, how mechanistically this regulation occurs, and whether particular factors regulate widespread chromatin state, remains largely unknown. Additionally, removal of major factors thought to function in regulation of chromatin accessibility, including H3K27ac, H3.3^2^, and prominent nucleosome remodelers^3^, has had only modest impacts on genome-wide chromatin state. Evidence indicates that the histone variant H2A.Z can promote binding of specific TFs, as well as transcriptional activators and repressors^4–7^, but the precise molecular mechanisms by which H2A.Z functions, and whether H2A.Z localization patterns influence genome-wide transcription factor (TF) binding events remains unknown.

The eukaryotic genome is almost completely wrapped by nucleosomes or bound by TFs^8,9^, with less than 5% of total genomic DNA sequence available in a non-nucleosomal context, regardless of cell type^8^. The vast majority of TFs can bind only at these accessible regions. Chromatin state varies considerably between cell types, but how chromatin state transitions occur and the role of epigenetic factors in driving these changes remains unclear^9^. The prevailing model for transitioning between chromatin states relies on passive competition between TFs and histones, but aside from pioneering factors, which bind at limited genomic locations^10,11^, the mechanisms by which the majority of genomic locations are converted from nucleosome bound to TF bound have not been identified.

Previously studies provide examples where H2A.Z localization can promote TF binding, including impacts on OCT4^6^ and FOXA2^7^ in stem cells. H2A.Z presence has also been correlated with DNaseI sensitivity at estrogen responsive enhancers in human breast cancer cells^4^. Whether H2A.Z nucleosomes are broadly favored for genome-wide TF binding, and the mechanisms by which these nucleosomes might harbor preferred TF binding sites are presently undetermined. Studies have shown that H2A.Z nucleosomes are more labile than canonical nucleosomes^12,13^. This concept gives rise to several possible ways in which DNA sequences at H2A.Z sites might be more exposed than the DNA of canonical nucleosomes. These include differences in the wrapping of DNA, differences in nucleosome turnover rates, and differences in chromatin stability at H2A.Z nucleosomes^14,15^. Additionally, increased nucleosome remodeling at H2A.Z sites, which has been observed in yeast and plants^7,16–18^, might allow for better access to DNA at H2A.Z nucleosomes compared with canonical nucleosome sites. Notably, it has not yet been established whether these characteristics of H2A.Z are important for mammalian TF binding.

H2A.Z has been most often studied in the context of transcriptional regulation. Studies have shown that decreased polymerase stalling correlates with the presence of H2A.Z at the ‘+1 nucleosome’ region of promoters^19^. In this manner, H2A.Z may promote gene activation by enabling RNA polymerase to overcome chromatin barriers at the earliest stages of gene activation^15,19–23^. However, separate studies have independently demonstrated that H2A.Z can be associated with gene silencing. Loss of H2A.Z causes particular genes to become active, and H2A.Z presence is necessary for establishment of the histone silencing mark H3K27me3 in stem cells^5,6,21,24^. Notably, much of the data supporting the dual role of H2A.Z in transcriptional regulation were acquired in stem cells, in the context of bivalent chromatin^25^, and it is presently unknown whether H2A.Z functions in direct silencing of transcription outside of this context. One additional possibility is that H2A.Z functions differently at distinct genomic locations. Genome-wide H2A.Z patterning is controlled by the H2A.Z incorporation complexes SRCAP and P400, and by the H2A.Z removal complex INO80^26–31^. Additionally, ANP32E has been identified as a chaperone protein that binds the C-terminus of H2A.Z in order to facilitate H2A.Z removal from chromatin^32^. Whereas deletion of H2A.Z is lethal, loss of ANP32E is compatible with viability in mice and zebrafish^33,34^. We and others previously found that ANP32E depletion leads to genomic H2A.Z reorganization, including regional H2A.Z accumulation^32,33,35^. Therefore, manipulation of *Anp32e* provides a means to assess locus specific function of H2A.Z.

Here, we use mouse fibroblasts lacking ANP32E to investigate how genomic H2A.Z organization regulates chromatin state, accessibility to DNA, and gene transcription. We find that modest changes in H2A.Z localization correspond with dramatic and widespread chromatin accessibility changes. Loss of ANP32E results in increased chromatin accessibility at sites where H2A.Z accumulates, but also at sites where high levels of H2A.Z in wild type (WT) remain stable in mutant cells. At thousands of H2A.Z marked promoters, we find that H2A.Z expands, spreading from the +1 nucleosome to the −1 nucleosome, and this expansion is associated with a dramatic increase in chromatin accessibility. In this regard, we determine that H2A.Z localization conforms to a hierarchy surrounding the transcription start site (TSS), and this hierarchy distinguishes strength of adjacent nucleosome positioning as well as TF motif accessibility. Finally, we find that ANP32E is necessary for activation of genes involved in cellular proliferation, and for silencing of genes involved in developmental differentiation. Thus, we establish that ANP32E controls genome-wide H2A.Z enrichment levels, and positioning of H2A.Z around the TSS in order to hone widespread gene expression levels.

## RESULTS

### H2A.Z resides at active genes and highly accessible chromatin regions

To begin investigating our hypothesis that H2A.Z contributes to gene activation and chromatin accessibility, we assessed two embryonic mouse cell types, mouse embryonic stem cells (MESCs) and mouse embryonic fibroblasts (MEFs). We found a high degree of overlap between H2A.Z and high chromatin accessibility at both promoter regions (**Fig 1A**) and H2A.Z sites genome-wide (**Fig S1A & 1B**). After partitioning the entire genome based on H2A.Z enrichment, we found that the highest levels of H2A.Z coincided with the highest degree of chromatin accessibility (**Fig 1C**), and the most accessible regions had the highest levels of H2A.Z (**Fig S1B**). This trend was even more apparent when we limited our analysis to promoters. Here, H2A.Z marked promoters were more highly expressed (**Fig 1D**), and the most accessible promoters were almost always marked by H2A.Z (**Fig 1E**). Additionally, gene expression was considerably higher for accessible promoters that were marked by H2A.Z, compared with promoters lacking either of these attributes (**Fig 1F**).

**Figure 1.**
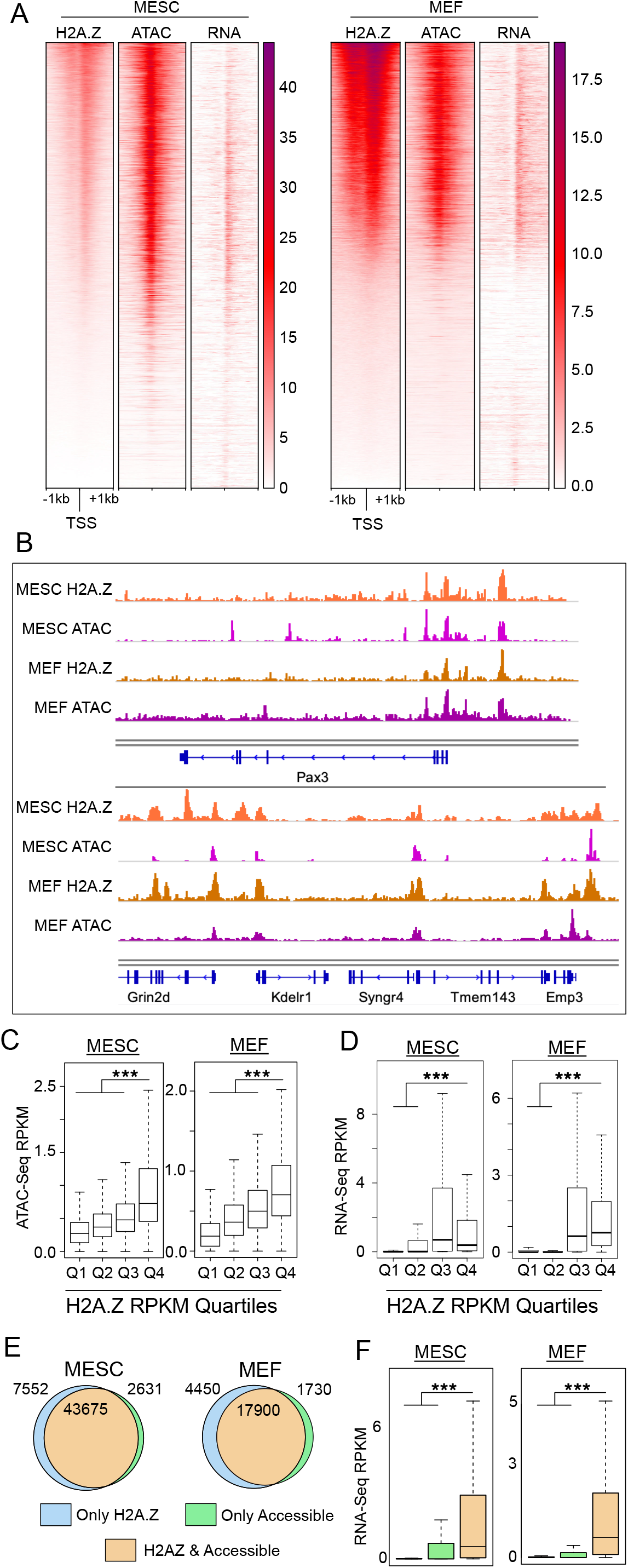
Genome-wide H2A.Z correlation with chromatin accessibility and gene expression across multiple cell types. (A) Heatmaps of normalized H2A.Z enrichment (ChIP-Seq – see methods), chromatin accessibility (ATAC-Seq – see methods) and RNA expression signals (RNA-Seq – see methods) at promoter regions (TSSs ± 1kb). Rows in each heatmap are ordered by decreasing H2A.Z signal, and only forward strand TSSs are shown. (B) A gallery of genome browser snapshots for representative genomic loci depicting the correlation of H2A.Z and chromatin accessibility (ATAC). Normalized reads counts are shown. (C) Boxplots of chromatin accessibility RPKM values for H2A.Z quartiles. A random sampling of regions (300bp, n=50,000) of mouse genome are used. Statistical significance is assessed with a two-sided Wilcoxon rank-sum test. (D) Boxplots of RNA expression RPKM values for H2A.Z quartiles at promoter regions (TSS ± 1kb). Statistical significance is assessed with a two-sided Wilcoxon rank-sum test. (E) Venn diagram showing overlap of H2A.Z peaks and chromatin accessibility peaks at promoter regions (TSS ± 1kb) for each cell type. The number of peaks in each category (H2A.Z occupied: light blue; accessible promoter regions: green, and both H2A.Z occupied and accessible regions: orange) are labeled. (F) Boxplots of RNA expression RPKM values for each category (same as panel D). Statistical significance is assessed with a two-sided Wilcoxon rank-sum test.

As noted, prior studies demonstrated that H2A.Z is necessary for recruitment of epigenetic silencing factors, including the PRC2 complex^5,6,24^, but in our initial analyses, we found that H2A.Z was strongly correlated with gene activation. Thus, we sought to further investigate how H2A.Z and chromatin accessibility associate with the activating epigenetic marks H3K4me3 and H3K27ac, and with the silencing mark H3K27me3. Here we found that H2A.Z is associated with bivalent chromatin marks H3K27me3 and H3K4me3 in MESCs (**Fig S1C** – top), consistent with prior studies^5,6,20,21^, but in MEFs, H2A.Z and H3K27me3 occur mostly at separate genomic locations (**Fig S1D**). Consistent with the cell-type-specific overlap of H2A.Z and H3K27me3, we found that H2A.Z was more highly correlated with gene activation in MEFs than in MESCs (**Fig 1C** – bottom). We also found cell-type-specific localization differences for H3K27ac comparing MEF enhancers with MESC enhancers, but unlike our measurements of H3K27me3, there was considerable overlap of H3K27ac, H2A.Z, and chromatin accessibility, regardless of cell type (**Fig S1E and S1F**). These results indicate that H2A.Z is consistently correlated with chromatin accessibility, as well as with marks of gene activation H3K4me3 and H3K27ac, but H2A.Z is only associated with the gene silencing mark H3K27me3 in the context of bivalent chromatin, at a subset of locations in MESCs.

### Loss of ANP32E leads to a widespread increase in chromatin accessibility

Having found a strong association of H2A.Z with highly accessible chromatin, we next sought to functionally test whether increasing H2A.Z enrichment levels, and altering H2A.Z genomic locations, would cause corresponding increases in chromatin accessibility. In separate mouse, human, and zebrafish studies, loss of the histone chaperone ANP32E resulted in genomic H2A.Z mislocalization ^32,33,35^. Therefore, we chose to investigate how chromatin accessibility changed in response to H2A.Z localization changes in primary MEFs lacking ANP32E. Our initial analyses took place at the genome-wide level, where unbiased clustering revealed strong concordance of biological replicates (**Fig S2A**). Loss of ANP32E resulted in dramatic and widespread increases in chromatin accessibility across the genome (**Fig 2A** - left). In fact, we observed heightened chromatin accessibility at more than 80% of all differentially accessible sites (**Fig 2B**). We did not expect such sweeping chromatin accessibility increases in *Anp32e* null cells based on prior reports that ANP32E loss leads to ectopic H2A.Z incorporation at approximately four thousand new sites^32^. In contrast, we found that accessibility increased at more than forty thousand sites, including TSS regions (**Fig 2A** - middle) and H2A.Z marked loci (**Fig 2A** - right).

**Figure 2.**
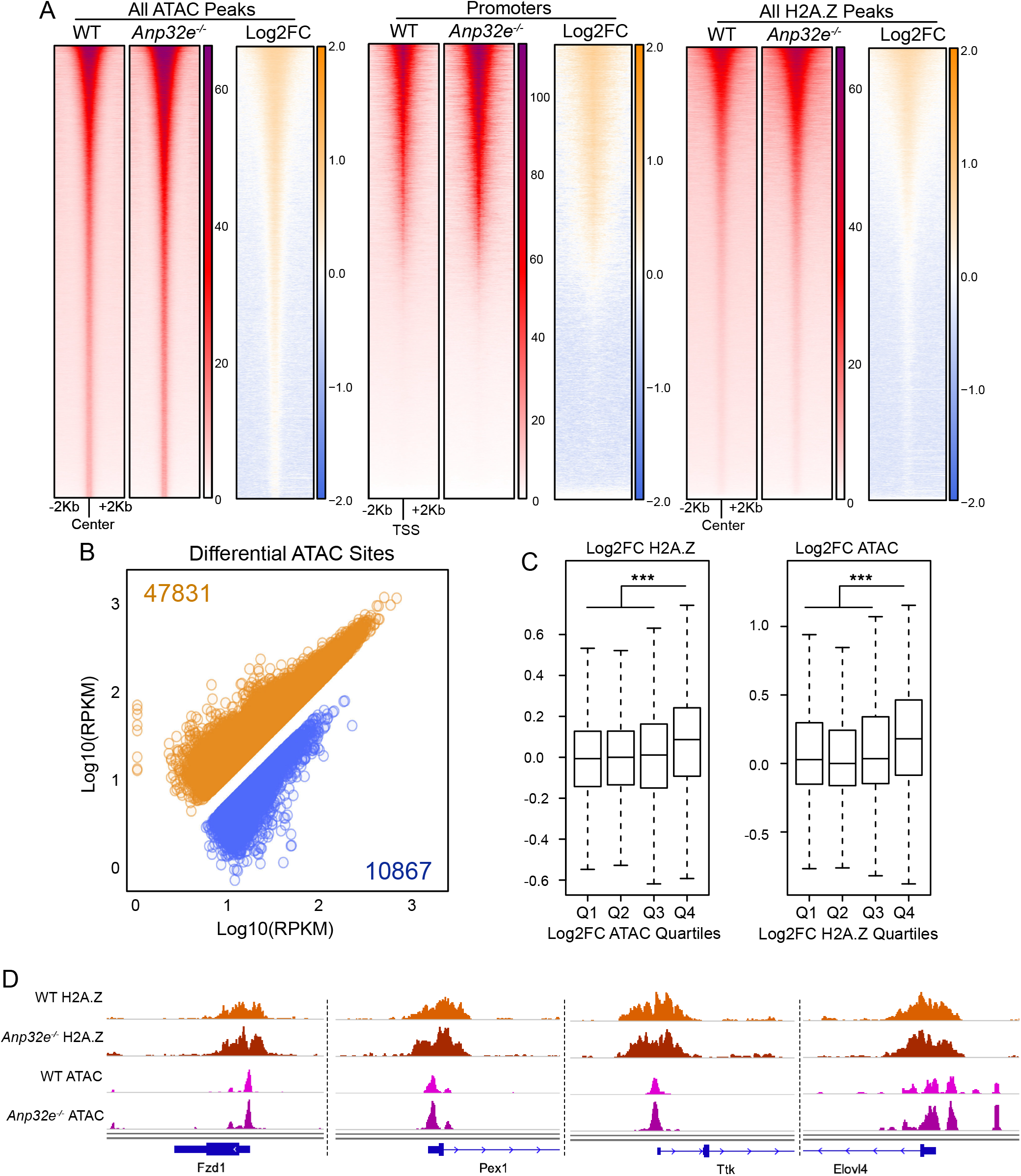
Genome-wide chromatin accessibility increases upon ANP32E loss. (A) Heatmaps of normalized chromatin accessibility reads (ATAC) and log2 fold change demonstrating a genome-wide increase in chromatin accessibility at all accessible sites (left panel), promoter regions (TSS ± 1kb, middle panel), and H2A.Z peaks (right panel) in *Anp32e*^−/−^ MEFs compared to WT. Union peaks of WT and *Anp32e*^−/−^ MEFs chromatin accessibility and H2A.Z enrichment are used for plotting. Ordering is by total ATAC signal. (B) Scatterplot of log10(RPKM+1) values illustrating increased chromatin accessibility. Increased ATAC sites in orange, and decreased ATAC sites in blue. Numbers of differentially accessible peaks are shown (see methods). (C) Boxplots of log2FC of H2A.Z enrichment (left) and chromatin accessibility (right) at TSS ± 1kb in WT and *Anp32e*^−/−^ MEFs. Statistical significance is assessed with a two-sided Wilcoxon rank-sum test. (D) Genome browser snapshots for representative genomic loci depicting increases in chromatin accessibility in *Anp32e*^−/−^ compared to WT MEFs.

If H2A.Z indeed functions to promote chromatin accessibility, then changes in H2A.Z levels might result in differing magnitudes of accessibility changes. We therefore investigated whether the observed chromatin accessibility increases in *Anp32e* null cells were accompanied by changes in H2A.Z levels. ANP32E loss led to an overall increase in H2A.Z enrichment, but we also found many sites where H2A.Z was either unchanged or reduced (**Fig S2B**). Regions with the greatest increase in H2A.Z were indeed those with the greatest increase in chromatin accessibility, and likewise, regions where chromatin become most accessible were also those where H2A.Z increased the most (**Fig 2C**). This trend was apparent throughout the genome (**Fig 2D**). However, we frequently found examples where heightened chromatin accessibility occurred at sites where H2A.Z levels remained unchanged (**Fig S2C**), and we also found modest increases in accessibility at sites where H2A.Z was somewhat reduced. The latter group represented the extreme minority of regions, and chromatin changes at these sites were much less significant than changes at all other sites (**Fig S2D**). These results indicate that changes in H2A.Z abundance do indeed confer changes in chromatin accessibility, but additional factors aside from H2A.Z enrichment might also contribute to accessibility changes.

### Positional hierarchy of H2A.Z at promoters coincides with differences in gene function

We next sought to identify additional H2A.Z specific chromatin changes that might account for the dramatic chromatin accessibility changes we observed. Several studies have reported H2A.Z enrichment at promoters flanking the TSS, including the first nucleosome located upstream of the TSS, termed the ‘−1 nucleosome’ position, and the first nucleosome located downstream of the TSS, termed the ‘+1 nucleosome’ position^21,36–38^. The 200-500bp regions located between these two nucleosome positions is often referred to as the nucleosome depleted region (NDR). In yeast, the H2A.Z installation enzyme Swr1 has been reported to localize at NDR sites to install H2A.Z at promoters^38–40^. These observations prompted us to investigate whether H2A.Z localization on either side of the TSS is impacted by ANP32E loss. Indeed, in the absence of ANP32E, H2A.Z accumulated mostly upstream of the TSS, near the −1 nucleosome position (**Fig 3A & S3A**). In order to better understand how H2A.Z might accumulate specifically at these sites, we categorized H2A.Z marked promoters based on relative changes in H2A.Z enrichment in response to ANP32E loss. Remarkably, at promoters where H2A.Z increased specifically at the −1 position, high levels of H2A.Z were maintained at the +1 nucleosome position in both mutant and WT conditions (**Fig 3B** – left **& S3B** – top). Alternatively, when H2A.Z accumulated specifically at the +1 nucleosome, these promoters generally lacked H2A.Z in WT MEFs (**Fig 3B** – right **& S3B** – bottom). Taken together, these results indicate that H2A.Z localization around the TSS adheres to a positional hierarchy, where *de novo* H2A.Z accumulates preferentially at the +1 nucleosome, and if H2A.Z is already present at the +1 nucleosome, additional H2A.Z accumulates at the −1 nucleosome.

**Figure 3:**
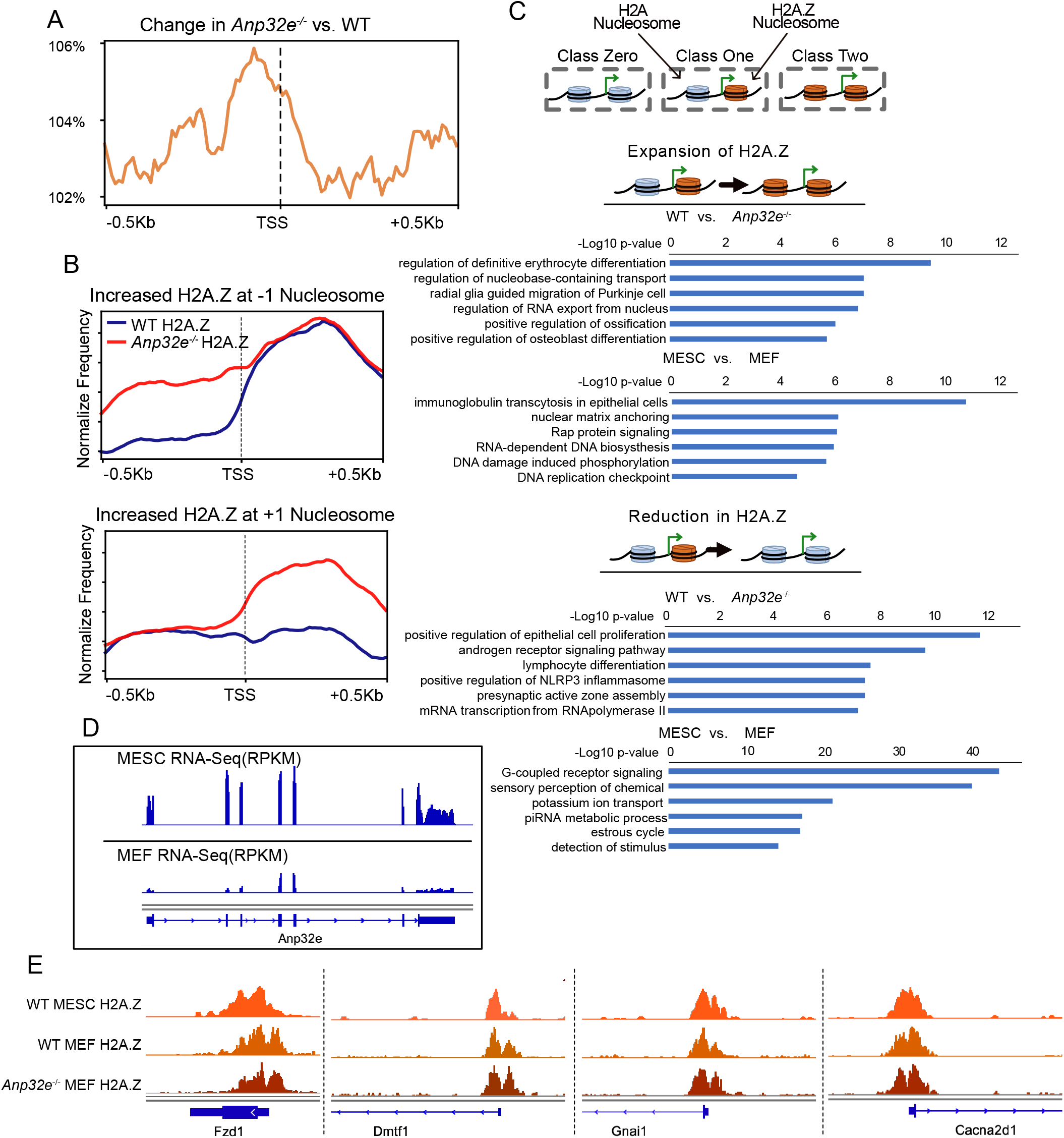
Changes in H2A.Z correspond with a positional hierarchy at genes with different function. (A) Change in H2A.Z showing an overall increase 5-prime of the TSS (indicated by a dotted line). Y-axis value represents calculated changes in H2A.Z levels comparing *Anp32e*^−/−^ to WT MEFs. Only forward strand TSSs are shown in the plot. (B) Aggregate H2A.Z enrichment plots in wildtype (blue) and *Anp32e*^−/−^ (red) at defined promoter categories. Regions where H2A.Z increased at −1 nucleosome are in the top panel, and regions where H2A.Z increased at +1 nucleosome are in the bottom panel. Only forward strand TSSs are shown in the plot. (See details in Methods). (C) Enriched gene ontology terms for promoter regions based on how H2A.Z classes change. Classes are depicted in Figure S3C. Locations that gain H2A.Z from Class One to Class Two (comparing either WT MEFs to *Anp32e*^−/−^ MEFs or WT MESCs to WT MEFs) are above, and locations that loss H2A.Z, going from Class One to Class Zero are below. (D) A genome browser snapshot of normalized RNA expression at the *Anp32e* locus showing higher *Anp32e* expression level in MESCs compared to MEFs. (E) Genome browser snapshot of H2A.Z signal in MESCs, MEFs, and *Anp32e*^−/−^ MEFs. Enrichment is RPKM.

We next asked whether ANP32E regulates H2A.Z localization at specific classes of gene promoters according to this positional hierarchy. To address this, we classified promoters based on H2A.Z localization changes in *Anp32e*^−/−^ MEFs relative to WT, and then assessed whether functionally distinct gene sets independently segregated (**Fig S3C**). Promoters lacking H2A.Z were defined as “Class Zero”, promoters with H2A.Z at only the +1 position were defined as “Class One,” and promoters with H2A.Z at both TSS flanking positions were defined as “Class Two”. Indeed, gene ontology (GO) analysis revealed that expansion of H2A.Z to the −1 nucleosome (going from Class One to Class Two) in *Anp32e* null MEFs occurred at genes associated with developmental differentiation (**Fig 3C** – top). Reduction of H2A.Z (going from Class One to Class Zero) occurred at several gene classes, with the most significant GO term being “positive regulation of epithelial cell proliferation” (**Fig 3C** – bottom). These results support a role for ANP32E in restricting H2A.Z expansion at promoters of genes involved in distinct biological processes. Based on our GO results for *Anp32e* null cells, we next wondered whether similar transitions between H2A.Z localization hierarchy classes at promoters occur during normal cellular differentiation. As a proxy for differentiation, we compared WT MESCs with WT MEFs, and again parsed promoters into defined H2A.Z positioning classes. Here we observed dramatic differences in H2A.Z positioning comparing MEFs with MESC (**Fig S3C** – bottom right), with a preference for H2A.Z expansions in MEFs as compared with MESCs. MESC Class One promoters became Class Two in MEFs just as frequently as they remained Class One (~40% of H2A.Z marked promoters). Similar to the ANP32E dependent H2A.Z changes we observed, distinct GO terms related to cell signaling (among other terms) were identified for genes where promoter H2A.Z localization changed (**Fig 3C**). One interesting possibility is that ANP32E levels and H2A.Z positioning are important for defining cellular function during differentiation and potentially for regulating H2A.Z positioning during cellular transitions. Consistent with this possibility, *Anp32e* mRNA is considerably lower in MEFs compared with MESCs (**Fig 3D**), and H2A.Z is much more restricted to the +1 nucleosome position in MESCs compared with MEFs (**Fig 1A & 3E)**.

### H2A.Z hierarchy changes correspond with accessibility and nucleosome positioning changes

Having found that ANP32E loss impacts H2A.Z localization patterns at gene promoters, we next asked whether these changes confer chromatin state changes surrounding the TSS. H2A.Z may promote increased accessibility through higher nucleosome turnover or increased lability. In this scenario, we expect that ANP32E loss would lead to reduced nucleosome positioning at H2A.Z sites. Alternatively, if H2A.Z promotes accessibility through nucleosome remodeling, we might expect that loss of ANP32E would lead to improved nucleosome positioning. To investigate this, we assessed nucleosome positioning by NucleoATAC-Seq^41^ and chromatin accessibility by ATAC-Seq^42^ in MEFS lacking ANP32E. We again categorized H2A.Z marked promoters based on H2A.Z changes at the −1 nucleosome, or the +1 nucleosome. Where H2A.Z increased at either of the TSS flanking nucleosome positions we observed both an increase in nucleosome positioning (**Fig 4A & S4A**), and an increase in accessibility over the TSS (**Fig 4B & S4B**), suggesting that nucleosome remodeling might underlie the observed accessibility changes (examples in **Fig 4C**). Recent reports from yeast indicate that the RSC nucleosome remodeler complex preferentially repositions H2A.Z containing nucleosomes^16^. If conserved in mammals, this mechanism might account for the observed relationship between H2A.Z expansion and nucleosome positioning in *Anp32e*^−/−^ cells. Strikingly, we found that nearly all H2A.Z marked promoters (**Fig 4D**) and enhancers (**Fig 4E**) in MEFs were enriched for SMARCA4, the closest mouse homolog to RSC^43,44^. At promoters, SMARCA4 was enriched mostly over the NDR, with H2A.Z containing nucleosomes enriched on both sides flanking SMARCA4 (**Fig 4F, S4C**). Taken together, these results are consistent with a nucleosome remodeling mediated mechanism where H2A.Z promotes increased nucleosome positioning flanking the TSS to stimulate chromatin accessibility at the NDR.

**Figure 4:**
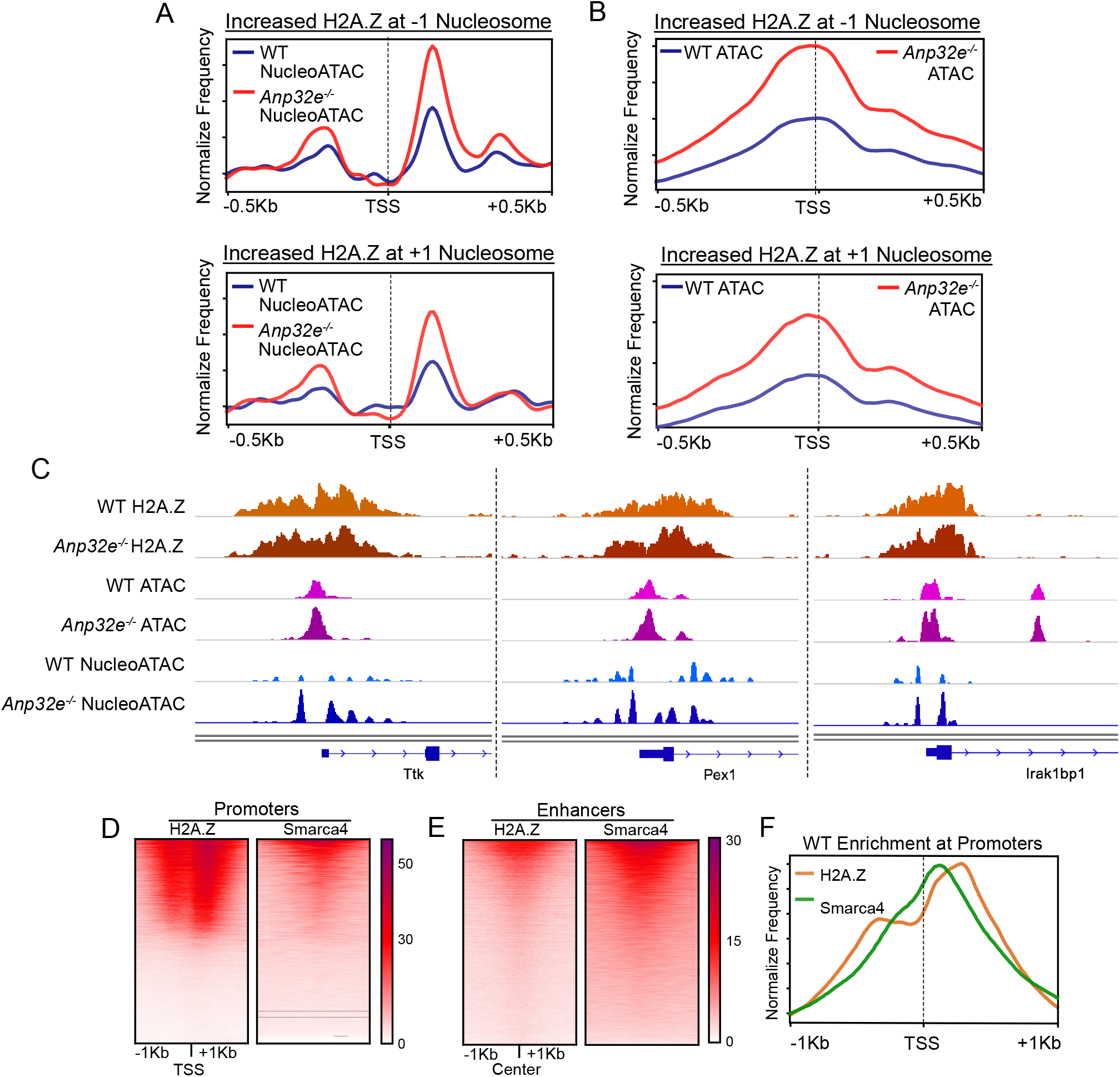
Changes in H2A.Z hierarchy correspond with nucleosome positioning and accessibility changes. (A) Aggregate NucleoATAC enrichment plots for WT and *Anp32e*^−/−^ signals at defined categories of promoters; those where H2A.Z increased at −1 nucleosome (top) and; those where H2A.Z increased at +1 nucleosome (bottom). Forward strand genes from each class are used to generate plots. (B) Aggregate total ATAC-Seq enrichment plots for WT and *Anp32e*^−/−^ signals at defined categories of promoters; those where H2A.Z increased at −1 nucleosome (top) and; those where H2A.Z increased at +1 nucleosome (bottom). Forward strand genes from each class are used to generate plots. (C) Genome browser snapshots for representative genomic loci depicting H2A.Z enrichment, increased chromatin accessibility (ATAC), and increased NucleoATAC signal for *Anp32e*^−/−^ compared to WT MEFs. (D) Heatmaps showing overlapping of H2A.Z enrichment and SMARCA4 enrichment at promoter regions (TSS ± 1kb). (E) Heatmaps showing overlapping of H2A.Z enrichment and SMARCA4 enrichment at putative enhancers. (F) Aggregate plots of H2A.Z enrichment and SMARCA4 enrichment at promoters (TSS ± 1kb). Forward strand genes are used to generate plots.

### TF-footprinting increases adjacent to better positioned H2A.Z nucleosomes

Our findings that loss of ANP32E leads to a greater degree of nucleosome positioning and increased chromatin accessibility at NDRs prompted us to investigate whether TFs might bind more effectively in response to ANP32E loss. As an indirect proxy for TF binding, we first assessed the abundance of non-nucleosomal chromatin fragments (NNFs). These NNFs were classified based on calculations from paired-end sequencing data as those fragments smaller than 150bp. These smaller fragments are known to accumulate as a result of TF binding rather than through nucleosome occupancy ^42^. We found that NNFs were more abundant in *Anp32e*^−/−^ cells than in WT MEFs (**Fig 5A**). Additionally, these NNFs were most abundant at Class Two promoters where H2A.Z marked both the +1 and −1 nucleosome position in WT MEFs (**Fig S5A**). In *Anp32e*^−/−^ MEFs, NNFs accumulated more at promoters where H2A.Z expanded from Class One to Class Two, compared with where H2A.Z was reduced to Class Zero (**Fig 5B**). This prompted us to further investigate whether genome-wide TF binding events were impacted by ANP32E loss. To identify putative TF binding events, we performed footprinting analysis using HINT-ATAC-Seq ^45^. Our assessment revealed footprinting over several known TF motifs, which are differentially protected in the *Anp32e*^−/−^ cells (**Fig S5B**). Consistent with increased NNFs at promoters, there were far more footprint sites genome-wide identified for TF motifs with positive HINT scores, than those with negative scores (**Fig 5C**), which is indicative of increased overall TF binding. Examples of motifs identified include those for NF-YB and SP1 (**Fig 5D & S5C**), which are known to physically interact and have roles in developmental differentiation^46–51^. Taken together, these results provide strong evidence that ANP32E loss causes expansion of H2A.Z at promoters, and this H2A.Z reorganization coincides with improved nucleosome positioning, and increased TF binding.

**Figure 5.**
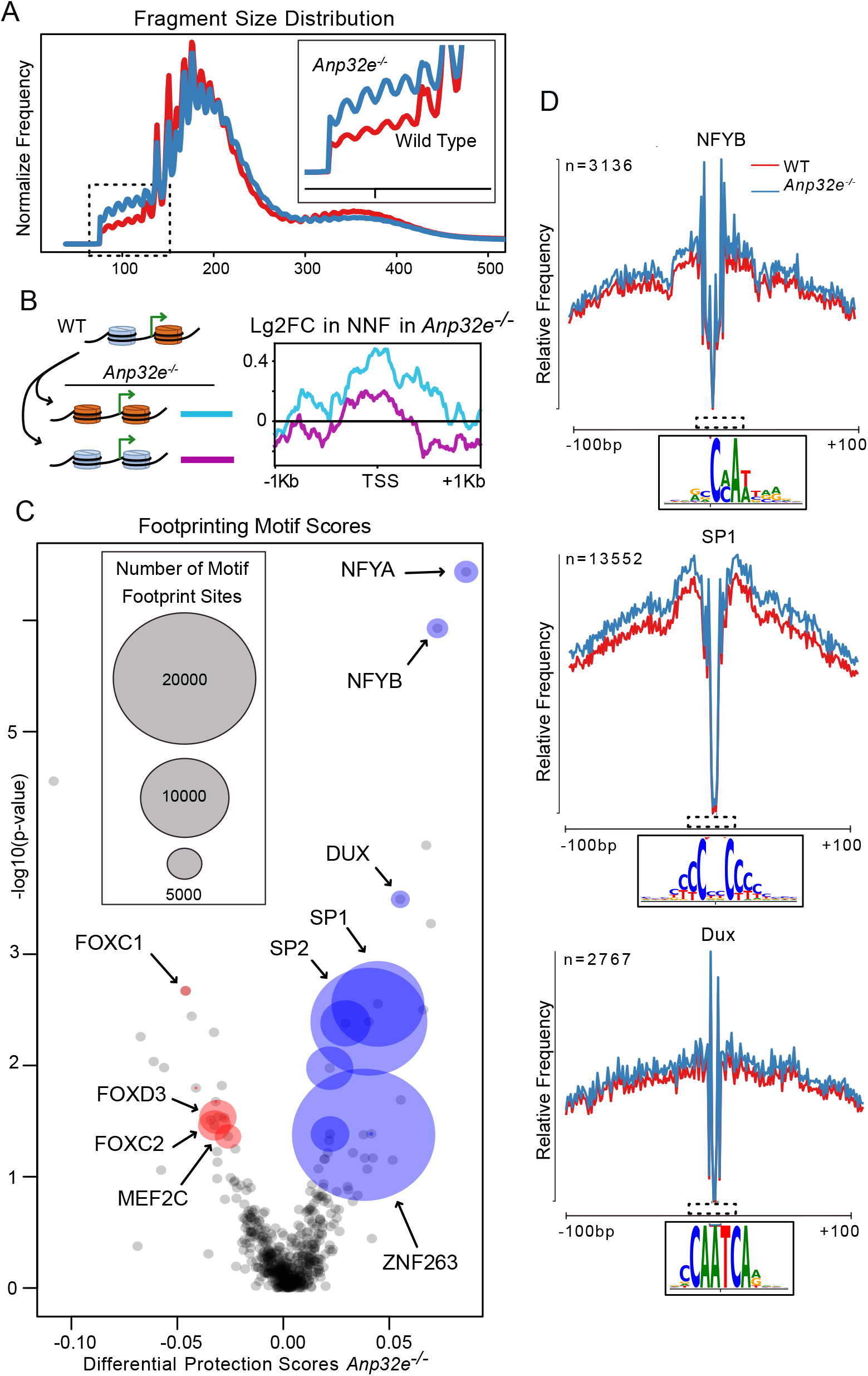
Hint-ATAC identifies differential transcription factor footprints in *Anp32*e depleted MEFs. (A) Distribution of fragment sizes in ATAC paired-end sequencing reads shows increased small fragment chromatin accessibility reads (< 150bp) for *Anp32e*^−/−^ (blue) compared to WT MEFs (red). Dashed box depicting small fragment reads is magnified. (B) Log2 fold change of small fragment chromatin accessibility reads (NNF, non-nucleosome fragments) for two clusters of genes; those that gain H2A.Z from ‘Class One’ to ‘Class Two’ promoter (blue) and those that lose H2A.Z from ‘Class One’ to ‘Class Zero’ promoter (purple) comparing WT MEFs *Anp32e*^−/−^ MEF. Log2FC of small fragment chromatin accessibility reads (<150bp) was generated by comparing *Anp32e* null chromatin accessibility to WT chromatin accessibility. Promoter classes are shown in Fig 3C. (C) Differential transcription factor footprints comparing *Anp32e*^−/−^ to WT MEF chromatin accessibility reads. Footprints of transcription factors are identified by HINT-ATAC using chromatin accessibility reads and narrowPeak files (see methods). Each point represents an individual TF, and the area of the point represents number of footprints identified for the corresponding TF. TFs with increased protection score in *Anp32e*^−/−^ are shown in blue, and TFs with decreased protection score in *Anp32e*^−/−^ are shown in red. Names of selected TFs with significant differential activity values are labelled. (D) Average cleavage profiles of NFYB, SP1, and Dux motifs identified by HINT-ATAC. A magnified view of the sequence motif (dashed box) for each TF is shown below plot. WT signal is in red and *Anp32e*^−/−^ is in blue.

### ANP32E loss leads to dysregulation of genes involved in differentiation

We expected that genome-wide transcriptional changes would occur as a consequence of the increased accessibility and TF footprinting that we observed. We therefore assessed genome-wide RNA expression levels (**Fig S6A**), and identified 1540 differentially expressed genes in the *Anp32e*^−/−^ cells (**Fig 6A**). Consistent with H2A.Z functioning as an activator, gene promoters with increased accessibility tended to be up-regulated, and promoters with reduced accessibility tended to be down-regulated (**Fig 6B**). Furthermore, we observed a more dramatic gene expression change when we restricted our analysis to genes in closest proximity to intergenic motifs identified by footprinting analysis (**Fig 6C**), indicating that the ANP32E mediated changes at intergenic regulatory regions, such as enhancers, may be a major contributor to the gene expression changes we observed. Similar to our investigation of promoters undergoing H2A.Z expansion, GO analysis revealed that genes critical for development were highly dysregulated (**Fig 6D**). Genes involved in pattern formation tended to be up-regulated, while genes involved in cell signaling tended to be down-regulated. Notably, GO terms associated with morphogenesis were present for both up- and down-regulated genes. Gene set enrichment analysis (GSEA) identified several developmentally important gene sets in which expression was significantly increased, and we found down-regulated genes to be associated with cell cycle and proliferation, including E2F target genes (**Fig 6E & S6B**). Examples of genes where accessibility and nucleosome positioning changes coincided with gene activation include *Crabp1* and *Cdkn1c* (**Fig S6C**). In sum, these results indicate that in addition to regulating H2A.Z localization and restricting genome-wide chromatin accessibility, ANP32E is a key regulator of developmental gene expression patterns.

**Figure 6.**
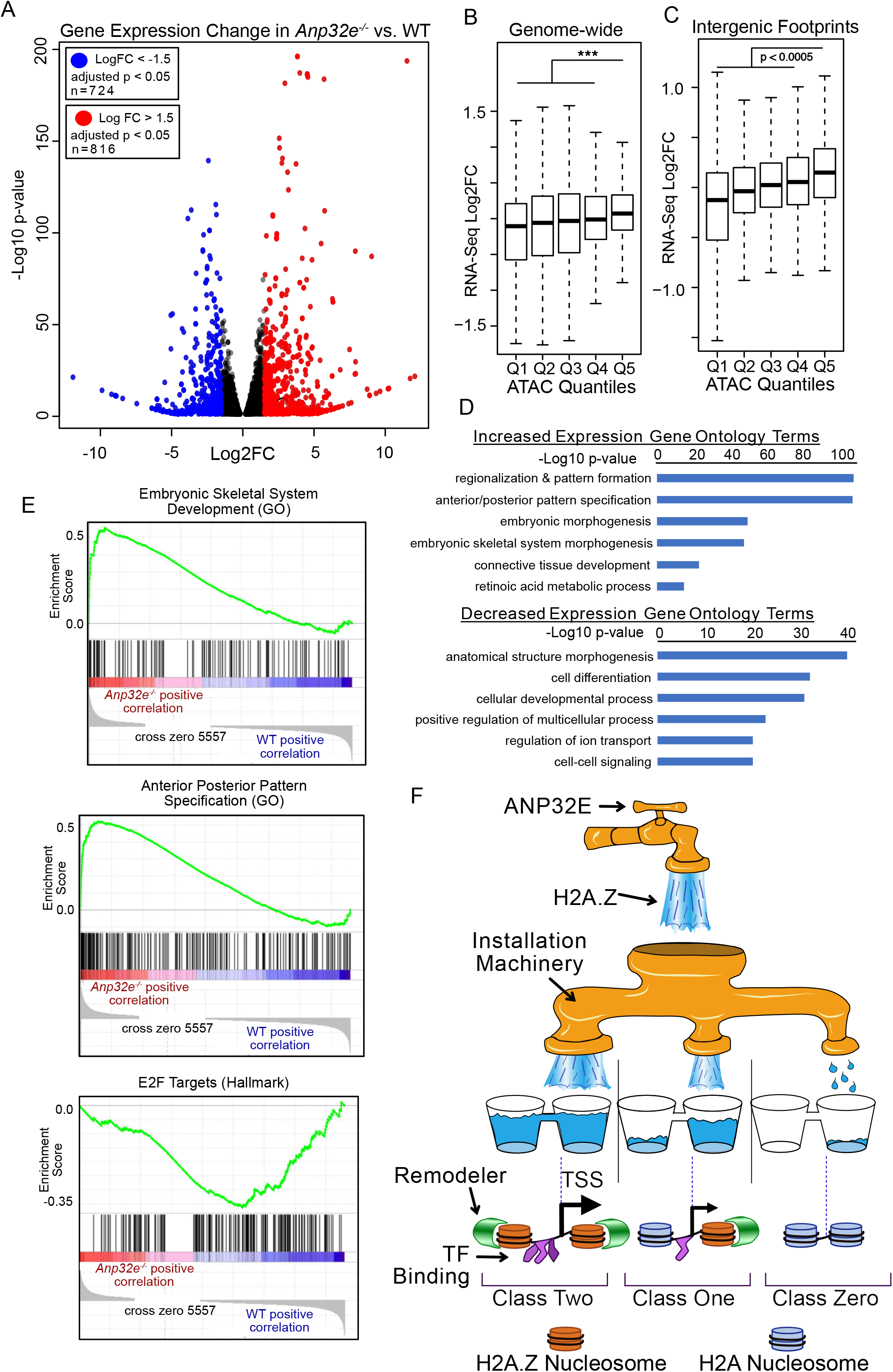
Differential gene expression profiles in *Anp32e*^−/−^ MEFs. (A) Volcano plot of differential gene expression comparing *Anp32e*^−/−^ to WT MEFs. Each point represents the average value of one transcript. Transcripts with significantly increased expression (log2 fold change > 1.5 and an adjusted p-value < 0.05) in *Anp32e*^−/−^ MEFs are shown in red, and those with significantly decreased expression (log2 fold change < −1.5 and an adjusted p-value < 0.05 are shown in blue. (B) Boxplots for log2 fold change of RNA expression at promoter regions (TSS ± 1kb) comparing *Anp32e*^−/−^ MEFs to WT MEFs. Quantiles are based on fold change in ATAC-Seq enrichment. Statistical significance is assessed with a two-sided t test. (C) Boxplots for log2 fold change of RNA expression at genes nearest to intergenic TF footprints comparing *Anp32e*^−/−^ MEFs to WT MEFs. Quantiles are based on fold change in ATAC-Seq enrichment. Statistical significance is assessed with a two-sided t test. (D) Enriched gene ontology terms for increased (left panel) and decreased (right panel) transcripts in *Anp32e*^−/−^ MEFs. (E) Gene set enrichment analysis of *Anp32e*^−/−^ compared to WT MEF RNA expression data revealed several gene sets and examples are shown. (F) A model for genomic H2A.Z localization and chromatin accessibility dynamics. ANP32E regulates H2A.Z abundance, and installation occurs in a targeted hierarchical manner. At locations where targeted installation is high, H2A.Z is installed at the +1 nucleosome position until it reaches maximum occupancy, and then H2A.Z is installed at the −1 nucleosome position. At locations where targeting is moderate, H2A.Z is installed only at the +1 nucleosome position, and where targeting is low, H2A.Z is absent. Where H2A.Z is most abundant, chromatin remodelers readily position nucleosomes flanking the nucleosome depleted region, TF binding increases, and neighboring gene expression increases.

## DISCUSSION

Here we demonstrate that ANP32E restricts genome-wide chromatin accessibility through regulation of H2A.Z accumulation and H2A.Z localization. In mouse fibroblasts lacking ANP32E, we found that heightened chromatin accessibility corresponded with increased H2A.Z, and with differences in H2A.Z positioning at promoters. We therefore define H2A.Z as a universal chromatin accessibility factor, and delineate three classes of promoters based on how H2A.Z is localized (**Fig 6F**). Class Zero promoters lack H2A.Z, have poorly positioned nucleosomes, and are largely inaccessible. Class One promoters have H2A.Z at the +1 nucleosome, have moderately positioned nucleosomes, and have intermediate levels of accessibility. Class Two promoters have H2A.Z at both the +1 and −1 nucleosome positions, have well positioned nucleosomes flanking the NDR, and have the highest degree of chromatin accessibility. Consistent with this hierarchical organization strategy, loss of ANP32E caused the greatest increase in short chromatin fragments, and the most TF motif protection at promoters where H2A.Z expanded to both TSS flanking positions. We found that this heightening of accessibility at TF binding motifs corresponded with increased gene expression. Thus, ANP32E abundance determines how genes are organized along the H2A.Z hierarchy. High levels of ANP32E drives promoters down the hierarchy to lower H2A.Z positioning classes, and low levels of ANP32E allows promoters to rise to higher H2A.Z classes. Taken together, these results indicate that through global regulation of H2A.Z, ANP32E restricts TF binding genome-wide, and in doing so, controls widespread transcriptional activation.

Cognizant of the various reports that H2A.Z has a dual function as an activator and a silencer^6,15,24,52^, at the onset of this study we expected to find two classes of H2A.Z marked genes, one class where H2A.Z expansion led to gene activation, and another where H2A.Z expansion led to silencing. However, when we assessed H2A.Z enrichment across two mouse embryonic cell types, we found that H2A.Z was frequently present at active genes, and H2A.Z was rarely present at silenced genes. Furthermore, H2A.Z marked gene promoters that gained more H2A.Z in the absence of ANP32E, and became more accessibility, tended to be up-regulated. Based on these results, we speculate that H2A.Z functions primarily as an activator. This view of H2A.Z aligns well with the vast majority of studies focused on H2A.Z transcriptional function, including early *Tetrahymena* studies describing H2A.Z presence in the transcriptionally active macronucleus and H2A.Z absence in the inactive micronucleus^53^, and studies describing H2A.Z as a semi-permissible or “labile” barrier to polymerase during gene activation^12,19,54,55^. To reconcile our findings with descriptions of H2A.Z functioning in gene silencing^5,6,24^, we favor a model where H2A.Z promotes activation in differentiated cells, but contributes to both activation and repression in pluripotent cells, perhaps through unknown cell-type specific mediators. Indeed, we did find evidence that H2A.Z corresponds with the silencing epigenetic mark H3K27me3 in mouse stem cells, but not in the other mammalian cell types we investigated, suggesting that H2A.Z might function in gene silencing uniquely in stem cells. It will be important for future studies to test this concept and determine whether H2A.Z function may indeed be cell-type or cell-state specific.

Our demonstration that H2A.Z promotes increased TF binding agrees with prior reports describing how H2A.Z function in OCT4 and FOXA2 recruitment^6,7^. Our results are also consistent with studies describing how H2A.Z depletion leads to decreased nucleosome levels^6^ in MESCs. It is commonly proposed that unique characteristics of H2A.Z nucleosomes account for region-specific differences in chromatin accessibility. For example, H2A.Z containing nucleosomes may be more loosely wrapped than canonical nucleosomes^37^, and this might facilitate increased TF binding. Based on this model, TF binding might occur without the need to displace the underlying nucleosomes, and one might expect that chromatin at the TF bound regions would harbor more weakly positioned nucleosomes. However, in our study we found that H2A.Z sites corresponded with more strongly positioning nucleosomes. Additionally, we found that increased accessibility occurred adjacent to, but not directly overlapping with H2A.Z sites, and we found that SMARCA4, the mammalian homolog for RSC, binds adjacent to H2A.Z sites. Therefore, we favor a model in which H2A.Z containing nucleosomes are the preferred substrate for nucleosome remodelers, and this preference leads to increased chromatin accessibility. We propose that increased nucleosome remodeling at H2A.Z sites leads to increased nucleosome repositioning, which uncovers TF binding motifs and promotes increased gene transcription. This model is in agreement with studies of chromatin remodelers in yeast and plants, which demonstrate that ISWI ^17^, RSC ^16^, and BRM^18^ prefer to remodel H2A.Z containing nucleosome. Although this mechanism has not yet been demonstrated in animals, SMARCA4 and H2A.Z do physically interact in MESCs^7^. Additionally, increased nucleosome turnover might occur as a function of remodeling, perhaps allowing for better TF binding. In this scenario, H2A.Z installation would increase without increases in total residency, possibly due to H2A.Z saturation. This mechanism might account for why some of the H2A.Z enrichment changes we observed were less dramatic than the observed chromatin accessibility changes. Future studies of mammalian factors are required to test whether there is functional conservation in nucleosome remodeling at H2A.Z sites.

In the present study, we found that ANP32E loss led to H2A.Z accumulation mostly at the −1 nucleosome position relative to the TSS. Rather than invoking an ANP32E targeting mechanism, we favor a simpler explanation where ANP32E functions to prevent H2A.Z incorporation. We propose that H2A.Z installation occurs in a targeted and hierarchical manner (**Fig 6F**), where H2A.Z is installed primarily at the +1 nucleosome, and at sites where H2A.Z is already present, ancillary H2A.Z is installed at the −1 position. Therefore, at promoters where targeted (SRCAP-mediated) H2A.Z installation is high, H2A.Z resides at both TSS flanking positions, and at moderately targeted promoters, H2A.Z resides only at the +1 position. Based on this concept, ANP32E might sequester free H2A.Z to partially inhibit installation, leading to H2A.Z enrichment only at the +1 nucleosome, and not the −1 nucleosome position. Loss of ANP32E would therefore allow for more H2A.Z availability, and ancillary H2A.Z installation would occur at the −1 nucleosome at moderately targeted promoters. This model fits well with prior studies reporting higher levels of H2A.Z at the +1 nucleosome position in mammals^21,35,56^, as well as studies in Drosophila and Arabidopsis where homologs of H2A.Z are absent from the −1 nucleosome position^15,57,58^. Additional studies are necessary to determine how H2A.Z targeting occurs, and to identify the factors that direct H2A.Z accumulation preferentially at the +1 nucleosome.

We found that loss of ANP32E in MEFs results in dramatic gene expression changes. Transcription increased at genes where small non-nucleosomal DNA fragments accumulated, and expression decreased where these fragments were lacking. These small non-nucleosomal fragments occurred mostly at promoters where H2A.Z levels increased and nucleosome positioning strengthened. Additionally, we identified motifs for known regulators of differentiation at these genomic locations, including SP1 and NF-Y. These results indicate that H2A.Z is a functional driver of heightened chromatin accessibility, leading to increased TF binding and activation of gene expression. Indeed, GSEA tests of our gene expression data indicate that when ANP32E is lacking, genes involved in developmental differentiation become activated and genes involved in cell cycle progression become silenced. Furthermore, when we performed independent gene ontology analysis on different classes of H2A.Z localization promoters, we again identified pathways associated with developmental differentiation for genes where H2A.Z increased, and pathways associated with cell cycle progression and proliferation for genes where H2A.Z decreased. The remarkable consistency between biological pathways identified for H2A.Z localization changes and transcriptional changes suggests that ANP32E might be a critical regulator of cell state transition. Our results imply that high levels of ANP32E may help to maintain proliferation and self-renewal, while low levels of ANP32E may permit cellular transitions and differentiation. This concept aligns well with recent studies of ANP32E in breast cancer^59^, where high *ANP32E* expression corresponded with activation of *E2F1*, more efficient G1/S transition, and cellular proliferation. Thus, it will be critical for future studies to investigate how levels of H2A.Z and ANP32E are balanced in a cell-type specific manner, and how these factors might regulate transcription factor binding, cell cycle progression, and cell state transitions.

Several reports suggest that the precise regulation of H2A.Z positioning might be critical for preventing cancer. Amplified expression of H2A.Z has been associated with poor prognosis for several cancer types, including glioma, melanoma, and breast cancer^60^. Studies of prostate cancer have found that H2A.Z positioning flanking *de novo* NDRs occurs in association with increased oncogene expression^61,62^, and in Madin-Darby Canine Kidney cells, differences in H2A.Z positioning flanking the TSS have been correlated with epithelial-to-mesenchymal transition^63^. Recent studies have also demonstrated that increased *ANP32E* expression in breast cancer is associated with tumor growth and proliferation^59^, perhaps occurring through the same mechanisms of H2A.Z regulation that we describe here. With consideration of these previously published results, our study suggests that precise control of ANP32E levels and H2A.Z positioning may be critical for preventing carcinogenesis. Thus, it will be important for future studies to investigate the mechanisms described here in to context of human diseases, including cancer.

## ACKNOWLEDGMENTS

We would like to thank Dr. Ali Hamiche, at the Institut de Génétique et de Biologie Moléculaire et Cellulaire, in Illkirch, France, for graciously donating MEFs. We thank Dr. Bradly Cairns at the Huntsman Cancer Institute, and Drs. Paula Vertino and Jeff Hayes at the University of Rochester Medical Center for helpful discussions. We thank the UR Genomics Research Center for library preparation and next generation sequencing. We specifically acknowledge Dr. Jeff Malik from the URMC-GRC for his assistance.

## AUTHOR CONTRIBUTIONS

KEM, FWM, and PJM designed experiments, interpreted data, and helped in manuscript preparation. FWM and PJM preformed the majority of bioinformatics analyses. KEM preformed all cell culture methods and wrote the initial manuscript. CEM assisted in biological sample preparation and assisted in bioinformatics analysis.

## COMPETING INTERESTS

None

## SEQUENCING DATA DEPOSITION

NIH GEO Datasets: GSE145705

## SUPPLEMENTAL FIGURE LEGENDS

**Supplemental Figure 1.**
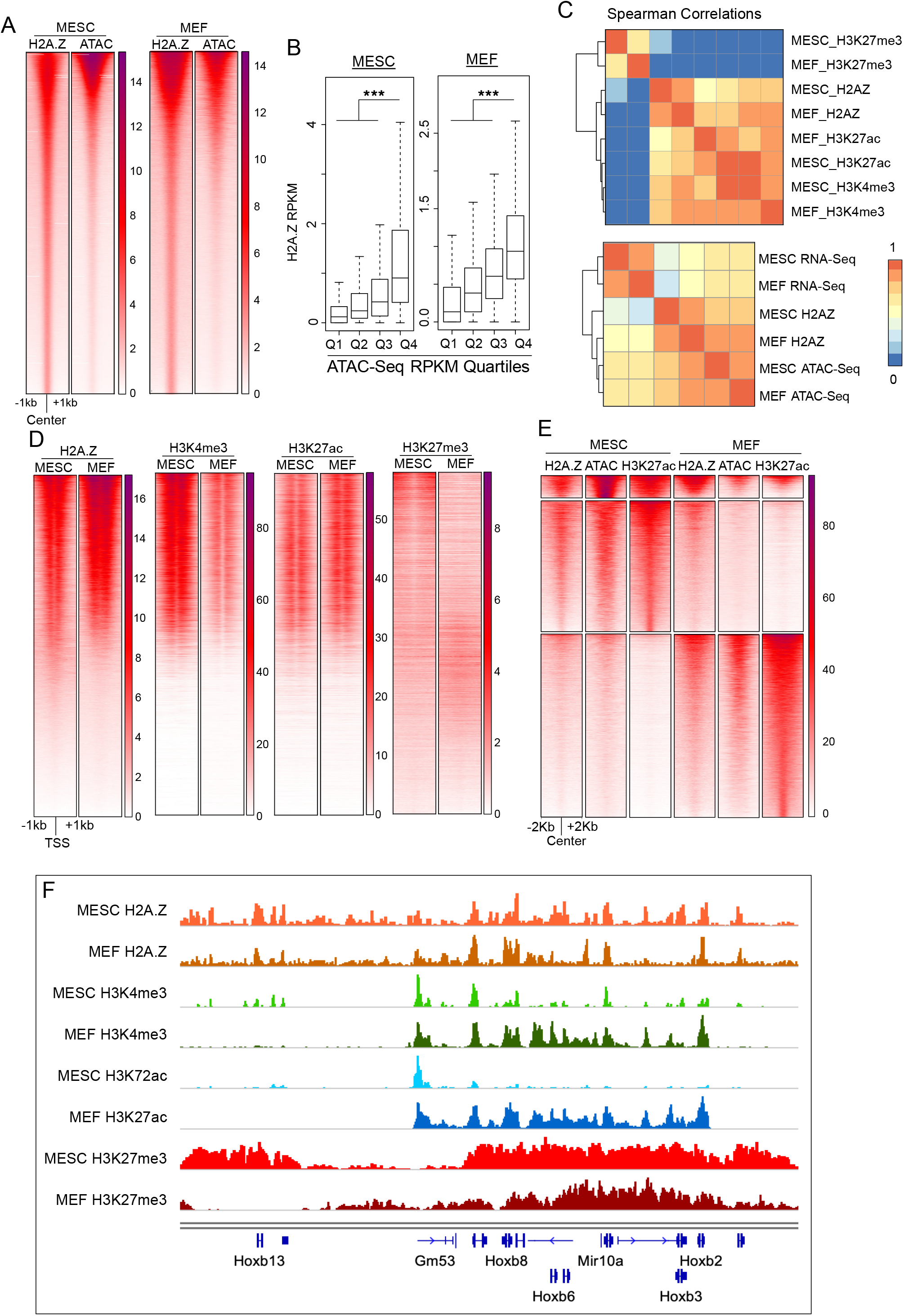
(A) Heatmaps of normalized H2A.Z enrichment and chromatin accessibility signals at H2A.Z peaks showing strong correlation of H2A.Z and chromatin accessibility. H2A.Z peak regions (peak center ± 1kb) are used to generate heatmaps. (B) Boxplots of H2A.Z enrichment RPKM values for chromatin accessibility quartiles showing that most accessible regions have higher H2A.Z levels. Randomly sampled regions (300bp, n=50,000) are used to for plotting. Statistical significance is assessed with a two-sided Wilcoxon rank-sum test. (C) Clustering of pairwise Spearman correlation values of several histone modifications and H2A.Z enrichment (top), as well as RNA, chromatin accessibility (ATAC) and H2A.Z enrichment (bottom) for MESCs and MEFs, displayed as heatmaps. (D) Heatmaps comparing enrichment signals of H2A.Z, H3K4me3, H3K27ac and H3K27me3 at promoter regions (TSSs ± 1kb) in MESCs and MEFs. All heatmaps were generated using the same ordered regions ranked by decreasing H2A.Z levels in MESCs and MEFs. (E) K-means clustering of heatmaps showing H2A.Z correlates with chromatin accessibility at putative enhancers. Putative enhancers are defined as a union of H3K27ac-occupied regions more than 2kb away from promoters in MESCs and MEFs. (F) Genome browser snapshots of the *Hoxb* locus showing H2A.Z enrichment and various epigenetic factors.

**Supplemental Figure 2.**
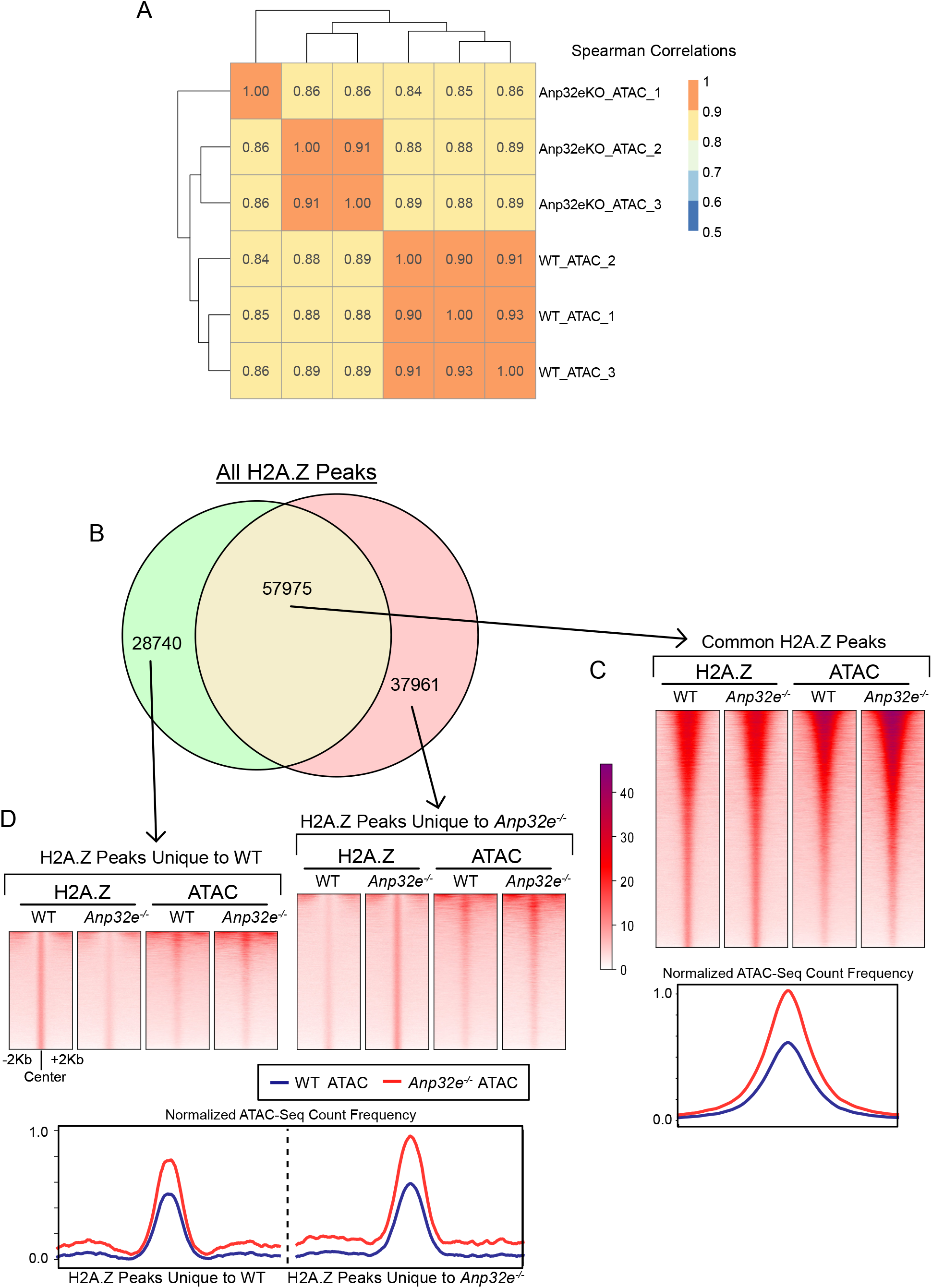
(A) A heatmap presentation of pairwise Spearman correlation values showing high correlation among chromatin accessibility replicates. (B) Venn diagram showing overlap of WT and *Anp32e*^−/−^ MEFs H2A.Z enrichment peaks. Numbers of H2A.Z peaks are shown in the plot. (C) Heatmaps of H2A.Z enrichment and chromatin accessibility (top), and aggregate plots of chromatin accessibility (bottom) show increased chromatin accessibility in *Anp32e*^−/−^ compared to WT MEFs at eH2A.Z peaks common to WT and *Anp32e*^−/−^ conditions (n=57,975). (D) Heatmaps of H2A.Z enrichment and chromatin accessibility (top), and aggregate plots of chromatin accessibility (bottom) show increased chromatin accessibility in *Anp32e*^−/−^ compared to WT MEFs at eH2A.Z peaks unique to WT (n=28,740, left) and unique to *Anp32e*^−/−^ (n=37,961, right).

**Supplemental Figure 3:**
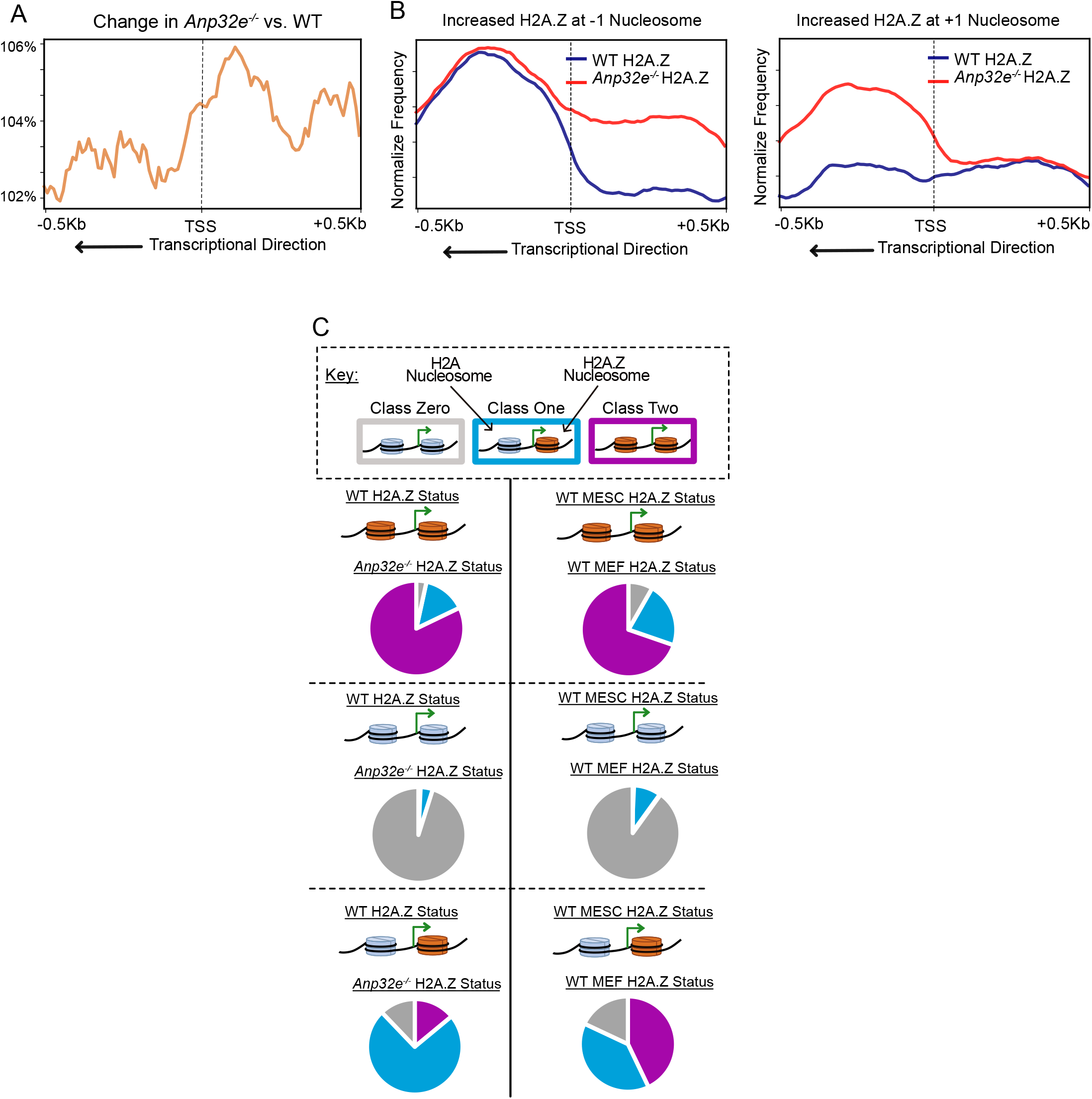
(A) Aggregate plot of percent change in H2A.Z and promoters depicting the direction of transcription. The dotted line indicates location of TSSs. (B) Aggregate plots of wildtype and *Anp32e*^−/−^ H2A.Z enrichment signals at defined categories of promoter regions: increased H2A.Z at −1 nucleosome (left panel), and increased H2A.Z at +1 nucleosome (right panel). Only reverse strand TSSs are shown in the plot. (C) Changes of H2A.Z status comparing WT MEFs to Anp32e^−/−^ MEFs (left) or WT MESCs (right). Initial H2A.Z positioning status (either in WT MEF or WT MESCs) is indicated by the nucleosome cartoon depicted within each panel, and how H2A.Z positioning changed is indicated by the accompanying pie chart. Color Key is included such that Class Zero promoters are grey, Class One promoters are blue, and Class Two promoters are purple.

**Supplemental Figure 4:**
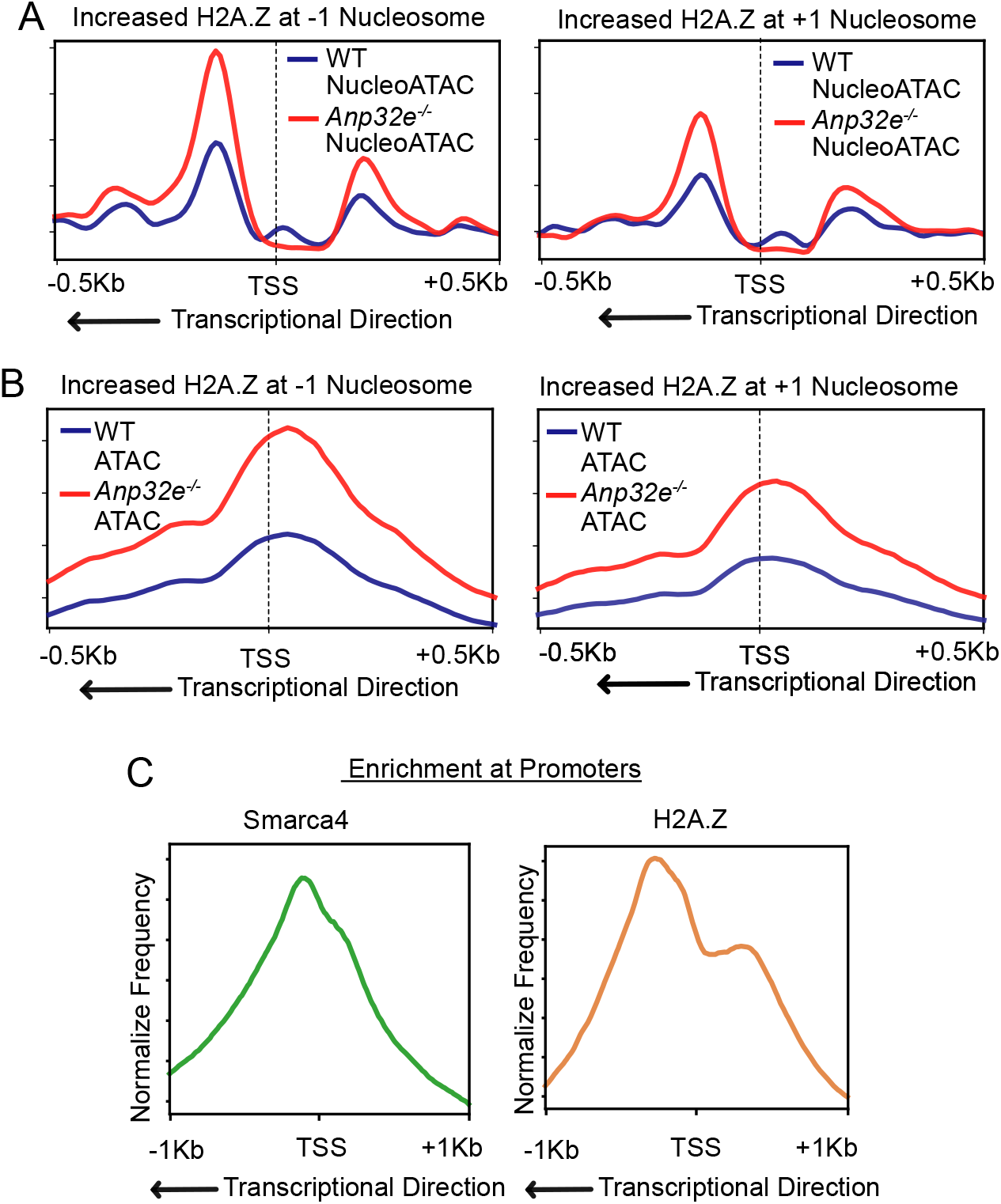
(A) Aggregate plots of WT and *Anp32e*^−/−^ NucleoATAC signals at two defined clusters of promoters. Promoters of reverse strand genes with increased H2A.Z at −1 nucleosome (left panel) and decreased H2A.Z at +1 nucleosome (right panel) are used to generate the plots. (B) Aggregate plots of WT and *Anp32e*^−/−^ chromatin accessibility reads at two defined clusters of promoters. Promoters of reverse strand genes with increased H2A.Z at −1 nucleosome (left panel) and decreased H2A.Z at +1 nucleosome (right panel) are used to generate the plots. (C) Aggregate plots of H2A.Z enrichment and SMARCA4 enrichment at promoters (TSS ± 1kb). Reverse strand genes are used to generate plots.

**Supplemental Figure 5:**
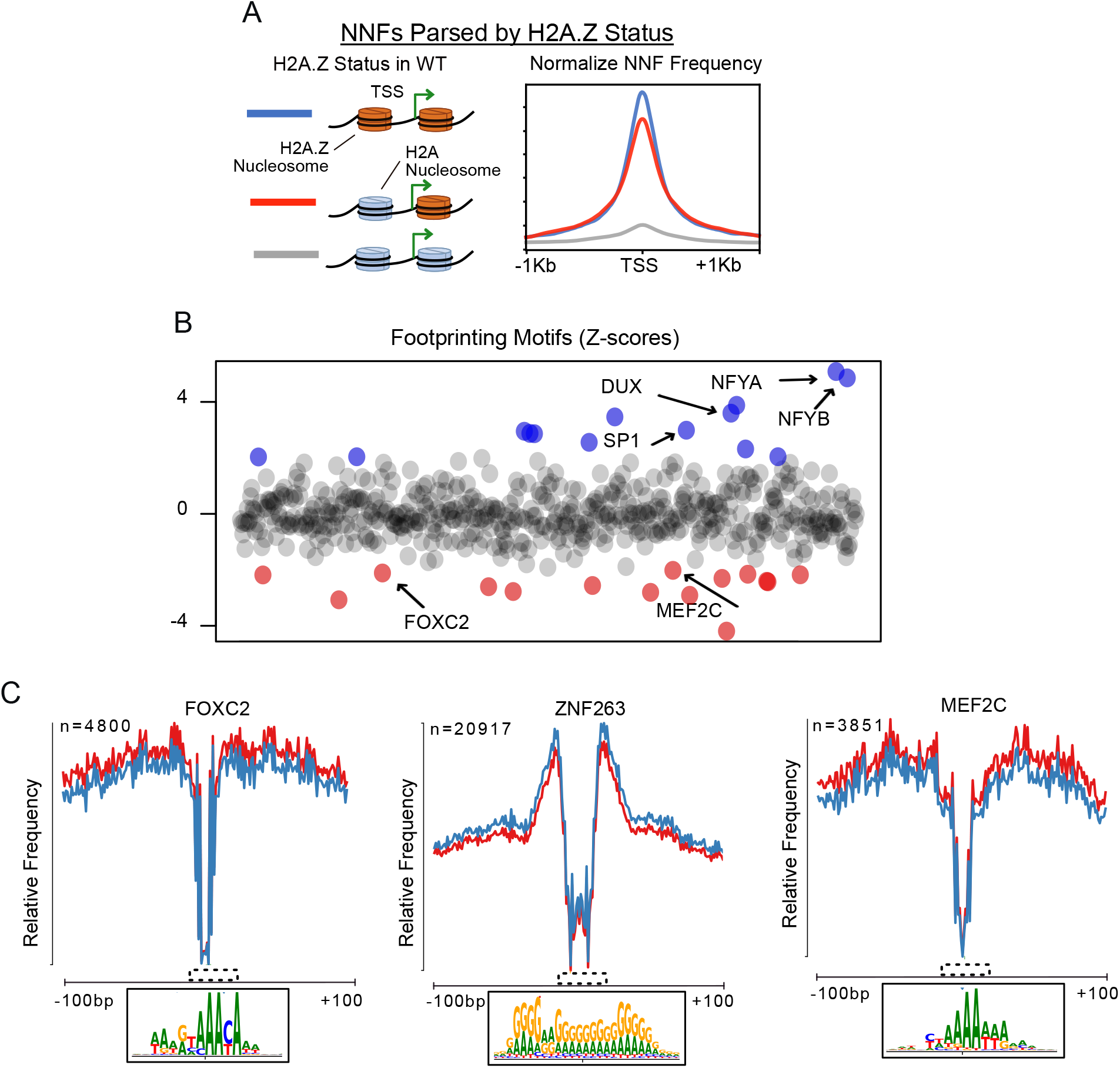
(A) Aggregate profiles of non-nucleosome fragments (NNF) of WT MEF chromatin accessibility reads for gene promoters parsed by H2A.Z status. Higher NNF distribution at ‘Class Two’ promoters (blue) than ‘Class One’ (red) and ‘Class Zero’ promoters (gray). (B) Scatter plot of transcription factors scores identified by HINT-ATAC footprinting analysis. Each point represents an individual TF. The y-axis represents the differences in TF activity comparing *Anp32e*^−/−^ to WT MEFs. TFs with significantly increased activity in *Anp32e*^−/−^ MEFs are shown in purple, and TFs with significantly reduced activity in *Anp32e*^−/−^ MEFs are shown in red. (C) Average cleavage profiles of FOXC2, ZNF263, and MEF2C motifs identified by HINT-ATAC. A magnified view of the sequence motif (dashed box) for each TF is shown below plot.

**Supplemental Figure 6:**
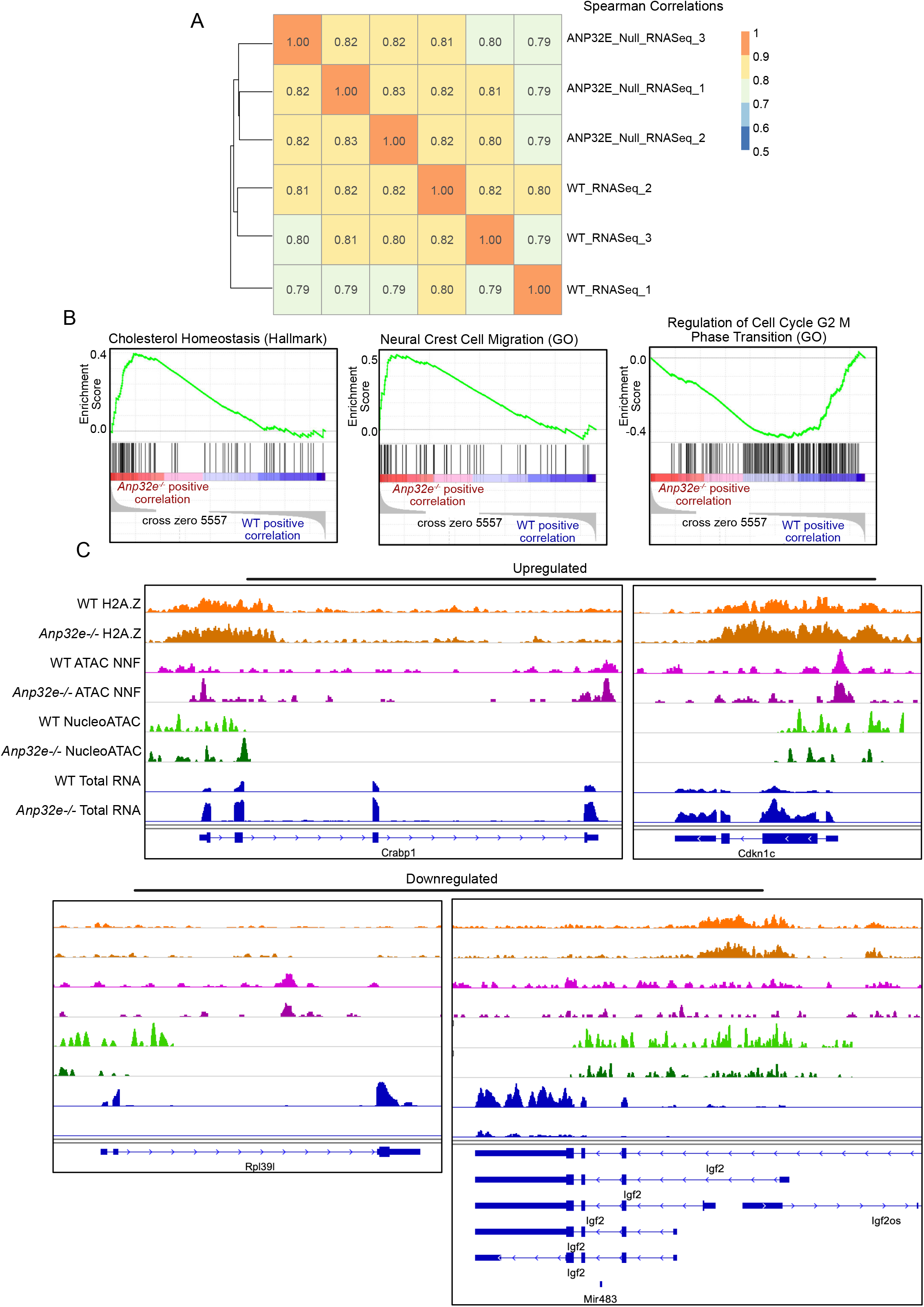
(A) Heatmap of pairwise Spearman correlation values showing high correlation among RNA sequencing replicates. (B) Gene set enrichment analysis of *Anp32e*^−/−^ compared to WT MEF RNA expression data revealed several gene sets and examples are shown. (C) Snapshots of genome browser views of H2A.Z enrichment, non-nucleosome fragment (NNF) of chromatin accessibility, NucleoATAC, and RNAseq of both wildtype and *Anp32e*^−/−^ MEFs.

## METHODS

### Cell Culture

Primary mouse embryonic fibroblasts were from WT and *Anp32e*^−/−^ embryos and graciously donated to us by Dr. Ali Hamiche, at the Institut de Génétique et de Biologie Moléculaire et Cellulaire, Cedex, France. MEFs were cultured in DMEM media (Thermo 11995073) containing 10% FBS and antibiotics. All assays were performed in parallel at passage numbers below 8. Approximately 200,000 cells were seeded on 10cm plates 1 day prior to RNA isolation or ATAC library preparation.

### Genomics Methods

Chromatin accessibility was measured using the Assay for Transposase-Accessible Chromatin combined with high throughput sequencing (ATAC-Seq), and gene expression levels were assessed using high throughput sequencing of RNA (RNA-Seq). Data were compared with data from studies which utilized Chromatin Immuno-Precipitation Sequencing (ChIP-Seq), but no new ChIP-Seq datasets were generated from this study. Data from numerous published studies were utilized ^24,32,64–66^. ATAC-Seq libraries were prepared using the Illumina Nextera DNA Flex Library Prep Kit (cat: 20018704) according to standard methods. RNA-Seq libraries were prepared using the TruSeq Stranded Total RNA Library Prep Kit (cat: 20020598), according to standard methods, and total RNA was purified using the Zymo Research Direct-zol RNA Miniprep kit (cat: R2051). Libraries were sequenced in 150bp paired-end sequencing mode on an Illumina NextSeq550 high-throughput DNA Sequencer. Sequencing took place at the University of Rochester Genome Research Center. Sequencing data deposited at NIH GEO Datasets: GSE145705.

### Bioinformatics Analysis

#### Alignment, normalization, and peak calling

For ChIP-seq and ATAC-seq, fastq files were aligned to mouse genome (Mm10) using Bowtie2, and then PCR duplicates were removed using Picard. Read count normalization was performed on alignment files to account for sequencing depth differences, and genome browser tracks in bigwig format were generated from merged replicates using deeptools bamCoverage. Peak calling was performed using MACS2 (setting for H2A.Z ChIP: -f BAMPE --SPMR --nomodel -B --broad; for ATAC-seq: -f BAMPE --SPMR --nomodel -B). Mitochondrial peaks and peaks overlapping blacklisted regions were filtered out for downstream analysis. Intersection of peaks were performed using bedtools intersect, and venn diagrams of peak numbers were generated using R package eulerr. Scoring differential ATAC-Seq peaks was based on absolute log2[(*Anp32e*^−/−^ ATAC reads+1)/(WT ATAC reads+1)] greater than 0.5, using the union peak set from WT and *Anp32e*^−/−^ MEFs.

#### Partitioning of promoters based on H2A.Z change

Normalized wildtype and *Anp32e*^−/−^ H2A.Z ChIP-seq bigwig file were used to generate average scores over 500bp window flanking both sides of TSS using deeptools. Then log2 H2A.Z fold change was calculated using H2A.Z average scores for both sides of TSS. Promoters with increased H2A.Z at −1 nucleosome are defined as log2FC values in top 25% at −1 nucleosome and bottom 75% at +1 nucleosome. Promoters with increased H2A.Z at +1 nucleosome are defined as log2FC values in top 25% at −1 nucleosome and bottom 75% at +1 nucleosome.

#### H2A.Z hierarchical positioning

Within deeptools, normalized wildtype H2A.Z ChIP-seq bigwig files was used to generate average scores from −600bp to −100bp (−1 nucleosome side) and 100bp-600bp (+1 nucleosome side) windows around TSS regions. The top 25% all H2A.Z average scores on both sides of TSSs was then used as the threshold value to partition promoters into different H2A.Z hierarchy categories (‘Class Zero’, ‘Class One’ and ‘Class Two’) in R. For example, when average H2A.Z enrichment was scored in the top 25% on both sides of the TSS, this was classified as ‘Class Two’ having H2A.Z at both −1 and +1 nucleosome position. To compare difference of H2A.Z hierarchical positioning between WT and *Anp32e*^−/−^ MEFs, threshold values established in WT were used. To compare MESCs with MEFs, threshold values established from MESC data were used. This partitioning of promoters occurred in R.

#### Transcription factor footprinting and nucleosome positioning

To identify transcription factors with differential footprints, narrowPeak files of wildtype and *Anp32e*^−/−^ ATAC-seq and duplicates-removed bam files were used as input files for Rgt-Hint-ATAC (-- atac-seq -- paired-end -- organism=mm10). Nucleosome positioning profiles were generated using duplicates-removed merged ATAC-seq bam files around promoter regions (TSS ± 1kb) with default setting of nucleoATAC. Genome browser tracks were generated converting nucleoatac_signal.smooth.bedgraph to bigwig format using bedGraphToBigWig.

#### Differential gene expression of RNA-seq data

RNA-sequencing fastq files were aligned to mouse genome (Mm10) using RNA-STAR. Read count normalization was performed on alignment files to account for sequencing depth differences, and genome browser tracks in bigwig format were generated using deeptools bamCoverage. A heatmap of spearman correlation using normalized reads across samples was generated using deeptools multibigwigsummary and R package pheatmap. Differential gene expression analysis was performed using Subread featureCounts and DESeq2 with default setting. Genes on X and Y chromosomes were excluded. Differentially increased genes were defined as log2[fold change] > 0.5 and adjusted p-value < 0.05, and differentially decreased genes were defined as log2[fold change] < −0.5 and adjusted p-value < 0.05.

#### Gene set enrichment analysis and Gene ontology

Gene set enrichment analysis of gene expression between wildtype and *Anp32e*^−/−^ MEFs was performed on reads count table generated by featureCounts using default setting of GSEA (broad institute). Gene ontology analysis of differentially increased and decreased genes was performed using PANTHER. Gene ontology of TSS from different H2A.Z hierarchical positioning was performed using GREAT^67^ with user input background datasets.

#### Plotting

Heatmaps and average aggregate plots were generated using deeptools plotHeatmap and plotProfile. Heatmaps of spearman correlation values were generated using R pheatmap, and boxplots and volcano plots are generated using standard R. Fonts and labels were adjusted with Affinity Designer.

### Statistical Methods

Comparisons between different quartiles of boxplot were performed using pairwise Wilcoxon rank sum test, with p-values adjusted using the “Hochberg” method, unless otherwise stated in the figure legends. Default statistical thresholds (p < 0.01) were applied as part of DESeq2 to identify differentially expressed genes in RNA-Seq analysis, unless otherwise stated. For correlational analysis, a Spearman correlation test was applied on complete datasets - regions where data was missing were excluded and R-values are indicated where applicable. Gene ontology analysis and gene set enrichment analysis were performed under default settings and p values are indicated in the plot. Hypergeometric test was applied to compare overlapping H2A.Z peaks and ATAC peaks, and p-values are indicated in the plot.

## Notes

### Competing Interest Statement

The authors have declared no competing interest.

